# Genomic Structural Variation Underlies Cell-Type-Specific Betacyanin Variegation in *Chenopodium quinoa*

**DOI:** 10.64898/2025.12.13.694152

**Authors:** Zheting Zhang, Yuwei Wang, Xiangwei Hu, Tiansheng Yu, Yaozu Feng, Jungao Zhang, Ting Zhang, Guojun Feng, Heng Zhang

## Abstract

The allotetraploid crop quinoa (*Chenopodium quinoa*) accumulates red/violet betacyanins, which function as vital stress-mitigating antioxidants. We investigated the genetic basis of red/green variegation observed in the aerial organs of the P0429 accession. We demonstrated that this color mosaic is primarily localized to the epidermal bladder cells (EBCs), with red EBCs accumulating ∼50-fold higher betacyanin levels than colorless EBCs. Cell-type-specific RNA-sequencing of EBCs identified the cytochrome P450 gene *Cqu0091301* (*CYP76AD*α) as the dominant and rate-limiting factor, exhibiting strong upregulation in red EBCs. This high pigmentation requires a specific structural variation in the P0429 accession: a ∼4-kb genomic insertion that restores the full functionality of *Cqu0091301*, which is otherwise truncated and non-functional in common reference genomes. Genomic analysis reveals that *Cqu0091301* is part of a *CYP76ADα*–*DODA* gene cluster. Notably, expression analysis revealed functional divergence between the quinoa subgenomes, with B-subgenome *CYP76ADα* genes highly dominant in EBCs, while A-subgenome homologs were preferentially expressed in other tissues. Our results establish a clear link between structural genomic variation and cell-type-specific betalain biosynthesis, providing molecular insight into pigment regulation and subgenome specialization in allotetraploid quinoa.

## Introduction

Variegation, defined as the occurrence of distinct color sectors within a single organ, is a widespread and important phenomenon in biology (Frank and Chitwood, 2016). In plants, this mosaic coloration typically results from cellular differences in chloroplast development, pigment biosynthesis, or dynamic epigenetic regulation among neighboring cell lineages (Frank and Chitwood, 2016). Classic examples include the *variegated 1* (*var1*) and *var2* mutants of *Arabidopsis thaliana*, where defective chloroplast biogenesis is caused by mutations in chloroplast protease genes (Yu et al., 2007). Furthermore, variegation can arise from genetic mosaics formed by spontaneous or induced mutations, as famously demonstrated by Barbara McClintock’s work on spotted maize kernels (McClintock, 1932). Transposon insertions and their epigenetic silencing have been linked to unstable pigment gene expression in maize, morning glory, and other species, resulting in cell-type-specific coloration patterns (Iida et al., 2004). Consequently, variegated phenotypes serve as powerful systems for studying the mechanisms governing gene expression stability, epigenetic control, and cell lineage determination.

Quinoa (*Chenopodium quinoa* Willd.) is an ancient pseudocereal renowned for its exceptional nutritional profile and strong tolerance to various abiotic stresses, particularly salinity (Alandia et al., 2020). As a facultative halophyte, quinoa does not require saline conditions for survival but is capable of completing the life cycle under 200 mM NaCl. A distinctive morphological feature of quinoa is the presence of specialized epidermal bladder cells (EBCs) covering most aerial organs (Shabala et al., 2014). EBCs are large, two-celled structures consisting of a stalk and a bladder cell, which was often considered the simplest form of salt glands (Dassanayake and Larkin, 2017; Bazihizina et al., 2022; Liu et al., 2024). EBCs have been reported to play critical roles in high salinity mitigation, osmotic regulation, and herbivore resistance (Kiani-Pouya et al., 2017; Kiani-Pouya et al., 2019; Bazihizina et al., 2022; Kiani-Pouya et al., 2022; Moog et al., 2023; Kobayashi and Fujita, 2024; Miranda-Apodaca et al., 2025).

The red/violet pigmentation observed in quinoa is attributed to betalains, a class of water-soluble nitrogenous pigments characteristic of the order Caryophyllales (Timoneda et al., 2019). Betalains functionally replace the more ubiquitous anthocyanins in most members of this order. Betalains are divided into two major groups – the red/violet colored betacyanins and the yellow colored betaxanthins – both derived from the amino acid L-tyrosine. The core betalain biosynthetic pathway involves three primary enzymatic steps: (1) hydroxylation of tyrosine to L-DOPA by a cytochrome P450 enzyme (CYP76AD); (2) ring cleavage of L-DOPA by DOPA 4,5-dioxygenase (DODA) to form betalamic acid; (3) subsequent spontaneous condensation reactions leading to betacyanin and betaxanthin formation (Timoneda et al., 2019). Cytochrome P450 enzymes of the CYP76AD family and DODAα isoforms play central roles in this pathway (Brockington et al., 2015; Polturak et al., 2016). Beyond their role as natural colorants, betalain pigments are potent antioxidants, with accumulation frequently linked to increased tolerance to environmental stressors like salinity, drought, and high light, primarily through the scavenging of reactive oxygen species and photoprotection.

Here, we investigate the genetic basis of a striking red and green mosaic variegated phenotype observed in the aerial organs of the quinoa accession P0429. We demonstrate that the color differences arise almost exclusively from the differential accumulation of betalains within the EBCs. Through cell-type-specific transcriptome analysis of isolated EBCs, we identified a key regulatory gene, *Cqu0091301*, a *CYP76ADα* homolog, whose expression strongly correlates with betacyanin accumulation in red EBCs. We further show that the reference genome annotation for this gene is incomplete; using genome resequencing, we discovered a ∼4-kb genomic insertion in P0429 that restores a complete, functional P450 domain. Comprehensive genomic analysis revealed that *Cqu0091301* is part of a multicopy *CYP76AD*α*-D*ODA gene cluster. Finally, we report an unbalanced contribution of the A and B subgenomes to betalain biosynthesis, with B-subgenome homologs showing preferential expression in EBCs and providing the dominant overall contribution across most pigmented organs. Our findings highlight a previously unrecognized role of structural genomic variation in controlling betalain biosynthesis and offer new insights into the cell-type-specific regulation of pigment formation and variegation in allotetraploid quinoa.

## Results

### Differential accumulation of betacyanins in EBCs underlies variegation in quinoa

We identified several quinoa accessions within our germplasm collection that exhibit a striking variegated phenotype, characterized by distinct red and green sectors on aerial organs, particularly the leaves (**Figure 1A; Figure S1A, B**). A cross-section of variegated leaves revealed that pigmentation is exclusively localized to the epidermal bladder cells (EBCs) in the red sectors, whereas the underlying leaf lamina tissue remained uniformly green (**Figure 1B**). Mechanical removal of the EBC layer resulted in the originally red sectors becoming visually indistinguishable from the green parts of the leaf (**Figure 1C**), indicating that variegation is caused by differential pigmentation in the EBC layer. The boundary between the two colored EBCs was sharp and developmentally stable, and occasionally aligned with the leaf vascular tissue (**Figure 1D**). Given previous reports of betalain accumulation in quinoa (Otterbach et al., 2021), we hypothesized that the variegated phenotype is caused by the differential accumulation of betacyanins in EBCs.

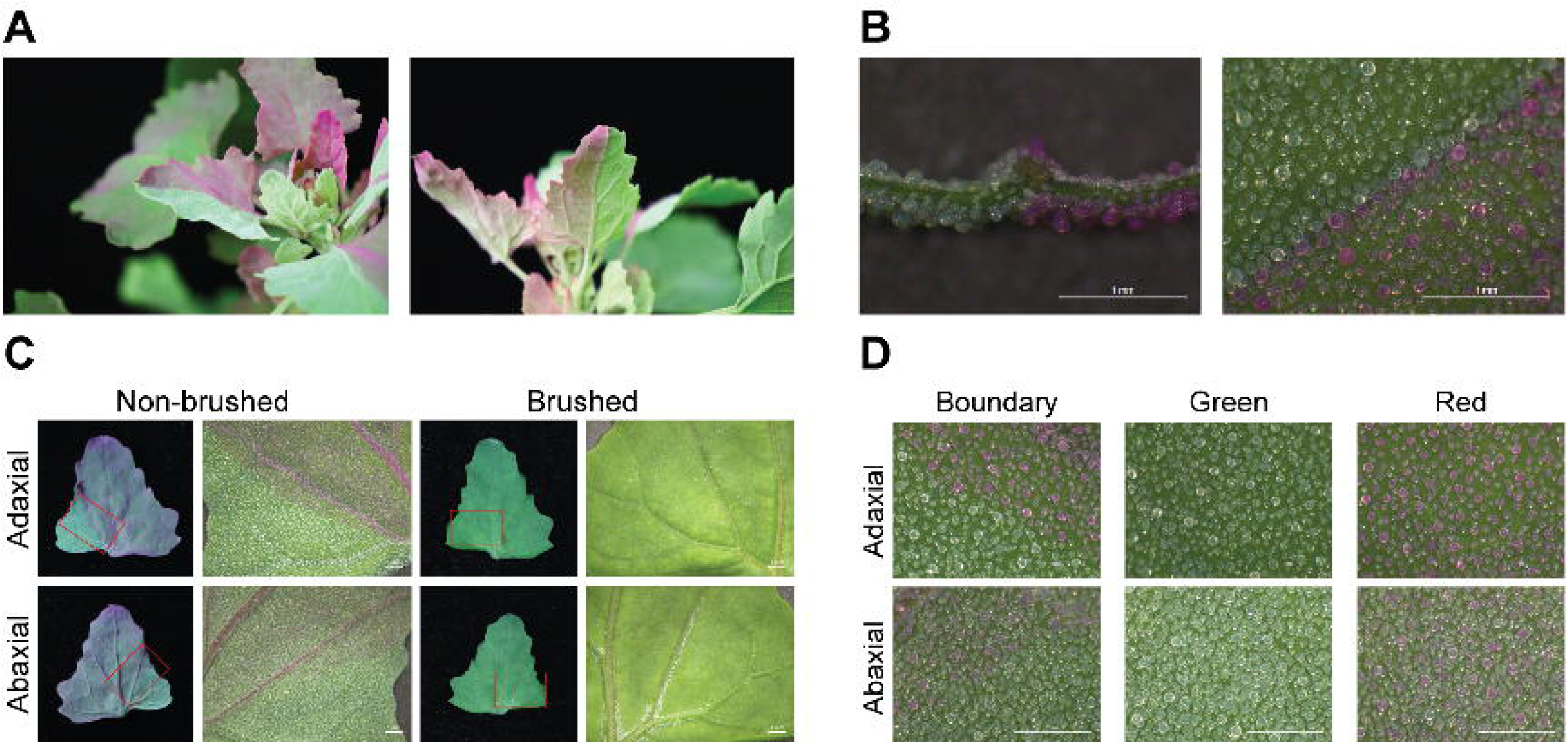

To quantify this difference, we measured betacyanin content in isolated red and colorless EBCs using a standard protocol (Stintzing et al., 2003). Water-soluble extracts from red EBCs exhibited a strong purple-red coloration, while extracts from colorless EBCs were nearly transparent (**Figure 2A**). Furthermore, the purple-red extract rapidly turned yellow upon the addition of sodium hydroxide, confirming the presence of betacyanins (**Figure 2B**). Spectrophotometric analysis at the betacyanin absorption maximum (538 nm) revealed that red EBCs contained 6.23 µg/g betacyanins, which is approximately 50-fold higher than the 0.12 µg/g measured in colorless EBCs from green sectors (**Figure 2C**). We also quantified betacyanins in leaf tissues after EBC removal. The red-sector lamina retained low levels (0.61 µg/g), while no detectable absorbance was measured in the green-sector lamina (**Figure 2C**). Crucially, the difference in betacyanin content between the red and green EBCs was an order of magnitude larger than the difference observed between the underlying lamina tissues. In contrast, no detectable absorbance was observed in the green sectors (**Figure 2C**). Moreover, we detected no significant differences in yellow betaxanthin levels (measured at 465 nm) between extracts from red and colorless EBCs (data not shown). Taken together, these results demonstrate that the leaf variegation phenotype is predominantly driven by the differential accumulation of betacyanins in the EBCs, with pigment differences in the underlying leaf lamina playing a minor role.

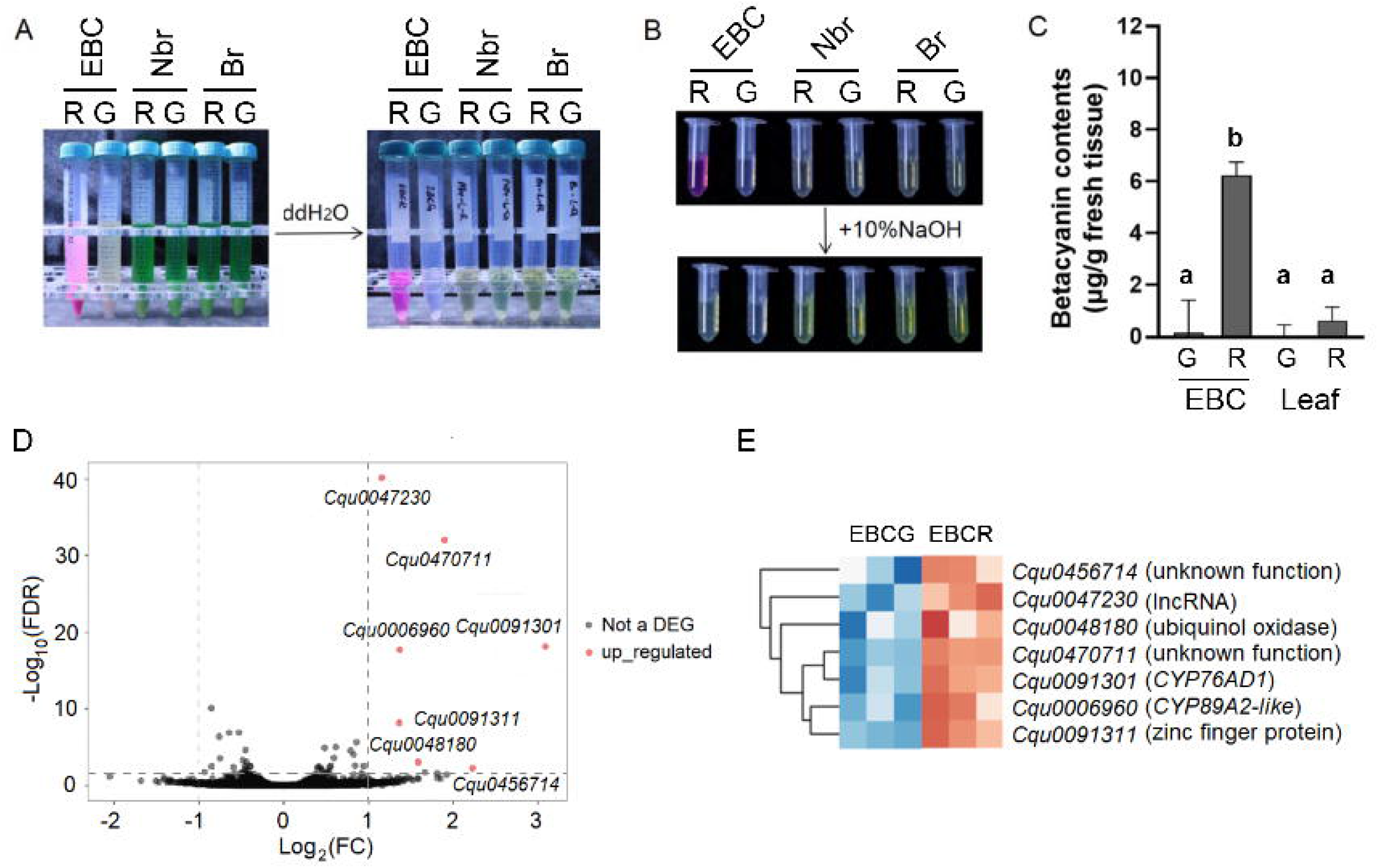

### Transcriptome analysis identifies genes associated with betacyanin accumulation

To identify the molecular mechanisms driving differential accumulation of betacyanins in epidermal bladder cells (EBCs), we performed RNA-seq on red and colorless EBCs isolated from the same variegated leaves of the accession P0429 (**Figure S1C**). Differential expression analysis yielded only seven differentially expressed genes, all of which were upregulated in red EBCs (**Figure 2D; Dataset S1**). The small number of DEGs was expected because EBCs were harvested from distinct sectors of the same leaf.

We next validated the expression level of seven DEGs using quantitative PCR (qPCR) in a new batch of EBC samples. Three genes (*Cqu0047230, Cqu0456714*, and *Cqu0470711*) were excluded from further analysis due to undetectable or unreliable expression (Ct values > 35). Of the remaining four genes (**Figure S1D**): (1) The P450 gene *Cqu0006960* (CYP89A family) showed low basal expression but was significantly upregulated in red EBCs. (2) *Cqu0091311* (zinc finger protein) was upregulated 4.5-fold. (3) *Cqu0048180* (ubiquinol oxidase) showed a statistically insignificant ∼2-fold increase. (4) *Cqu0091301* showed both the highest absolute mRNA level and the largest fold change compared to colorless EBCs. Notably, Significantly, *Cqu0091301* belongs to the CYP76ADα-type P450 family, which contains key components of the betalain biosynthetic pathway (Brockington et al., 2015). In contrast, *Cqu0006960* belongs to the CYP89A family, whose members are typically involved in leaf senescence (Mach, 2013). Given its high expression level, large fold change, and strong association with betalain synthesis, we prioritized *Cqu0091301* for further mechanistic analysis.

### *Structural variation of the* CYP76ADα *gene*

Inspection of the gene models of *Cqu0091301* in Cq_real_v1.5 and previous reference genomes (Jarvis et al., 2017; Zou et al., 2017) revealed a truncated second exon, resulting in the loss of a conserved P450 domain (**Figure 2A**). This structural anomaly suggested that the encoded enzyme was likely non-functional. However, RNA-seq data revealed a complex situation: (1) no reads aligned to the short second exon of *Cqu0091301* (**Figure S2A**); and (2) *Cqu0091280*, the closest homolog, showed a greater than 10-fold enrichment of mismatched (SNP-containing) reads mapping specifically over a portion of its second exon compared to other regions (**Figure S2B**). We hypothesized that these mismatched reads originated from an unannotated, highly homologous transcript in accession P0429 that was absent or incorrectly modeled in the reference genome. This was confirmed by *de novo* transcriptome assembly, which recovered a transcript that perfectly matched the annotated *Cqu0091301* mRNA but contained an extended 3’ region (data not shown). These findings collectively suggested that the P0429 genome contains a genomic insertion at this locus, resulting in a full-length, more complete *Cqu0091301* gene compared to the reference.

To precisely map this genomic variation, we performed whole-genome resequencing of P0429. Visual inspection of alignment reads in IGV revealed numerous soft-clipped sequences aligning downstream of the *Cqu0091301* locus (**Figure S3A**). We performed *de novo* assembly but were unable to reconstruct the full insertion. However, we identified a 2,273-bp contig that matched the flanking sequence of the insertion site on one end. Using targeted PCR with one primer on this contig and the other on *Cqu0091301*, we successfully amplified and sequenced the full 3,957-bp insertion missing in the reference genome but present in P0429 (**Figure 3A; Dataset S2**). Re-annotation based on this new sequence confirmed that *Cqu0091301* in P0429 contains a significantly longer second exon, restoring the complete, conserved P450 domain.

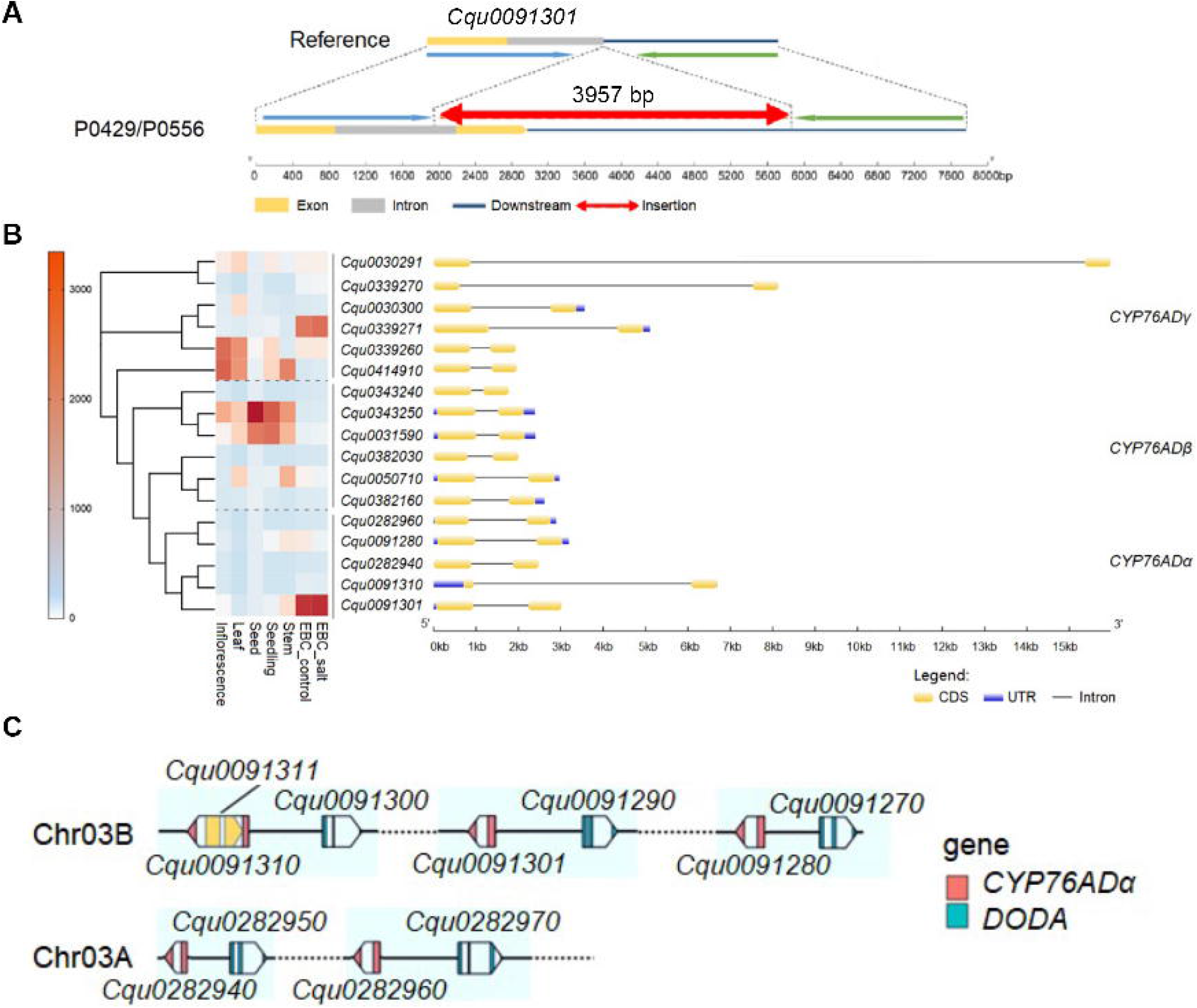

Re-analysis of the RNA-seq data using the corrected *Cqu0091301* gene model resulted in a more pronounced expression difference between red and colorless EBCs (**Figure S2C**), and, conversely, a reduction in the apparent expression difference for the homolog *Cqu0091280* (**Figure S2D**). The expression patterns of the remaining DEGs were unaffected.

To contextualize the role of *Cqu0091301*, we further examined the expression of other betalain biosynthetic genes, including *TyDC* (Tyrosine decarboxylase), *DODA, cDOPA5GT* (cDOPA glucosyltransferase), *B5GT* (betanidin 5-glucosyltransferase), and *B6GT* (betanidin 5-glucosyltransferase) in EBCs. Most of these genes showed extremely low mRNA levels in EBCs (often an order of magnitude lower) and did not exhibit differential upregulation in red EBCs (**Figure S4A**). QRT-PCR analysis corroborated these patterns (**Figure S4B**). The only genes consistently and markedly upregulated in red EBCs were the three CYP76ADα genes: *Cqu0091301, Cqu0091280*, and *Cqu0091310* (**Fig. S4B**). Taken together, these expression data strongly indicate that the upregulation of the CYP76ADα genes, particularly *Cqu0091301*, is the key regulatory step driving betacyanin synthesis in red EBCs.

To understand the genomic context of *Cqu0091301*, we identified 17 members of the *CYP76AD* family in the quinoa genome based on sequence homology. These genes are grouped into three subfamilies (α, β, and γ), each containing 5-6 members (**Figure 3B; Figure S5**). All the CYP76AD genes contain 2 exons, but their intron sizes are highly variable, ranging from 0.3 ∼ 14 kb (**Figure 3B**). All five *CYP76ADα* genes are physically linked and immediately adjacent to a *DODA* gene, forming *CYP76AD*α*–DODA* gene pairs located on two homologous chromosomes (**Figure 3C**).

### Unbalanced contribution of the AB subgenome to betalain biosynthesis

Beyond EBCs, the P0429 accession also exhibited distinct pigmentation patterns across various organs, including sharply segregated red and green sectors in the inflorescence (**Figure 4A–C**), red epidermal stripes on the mature stem (**Figure 4D, E**), and red pigmentation in the hypocotyl and cotyledons of seedlings (**Figure 4H, I**). We therefore investigated whether the *CYP76ADα* genes found to be critical in EBCs also regulate pigmentation in other tissues. Using qRT-PCR, we examined the mRNA level of the five *CYP76ADα* genes in green- and red-color sectors of seedlings, stems, and flowers. Among these homologs, the three B-subgenome copies *Cqu0091280, Cqu0091301*, and *Cqu0091310* showed a strong preferential expression in EBCs (**Figure 4J**). In contrast, the mRNAs of A-subgenome genes *Cqu0282960* and *Cqu0282940* were not detected in EBCs; *Cqu0282960* was preferentially expressed in seedlings, and *Cqu0282940* exhibited preferential expression in stems and flowers (**Figure 4K**). The expression of both genes was significantly up-regulated in the red tissues (**Figure 4K**). These results indicate that *CYP76ADα* genes from the A- and B-subgenomes have evolved distinct, non-overlapping tissue expression preferences.

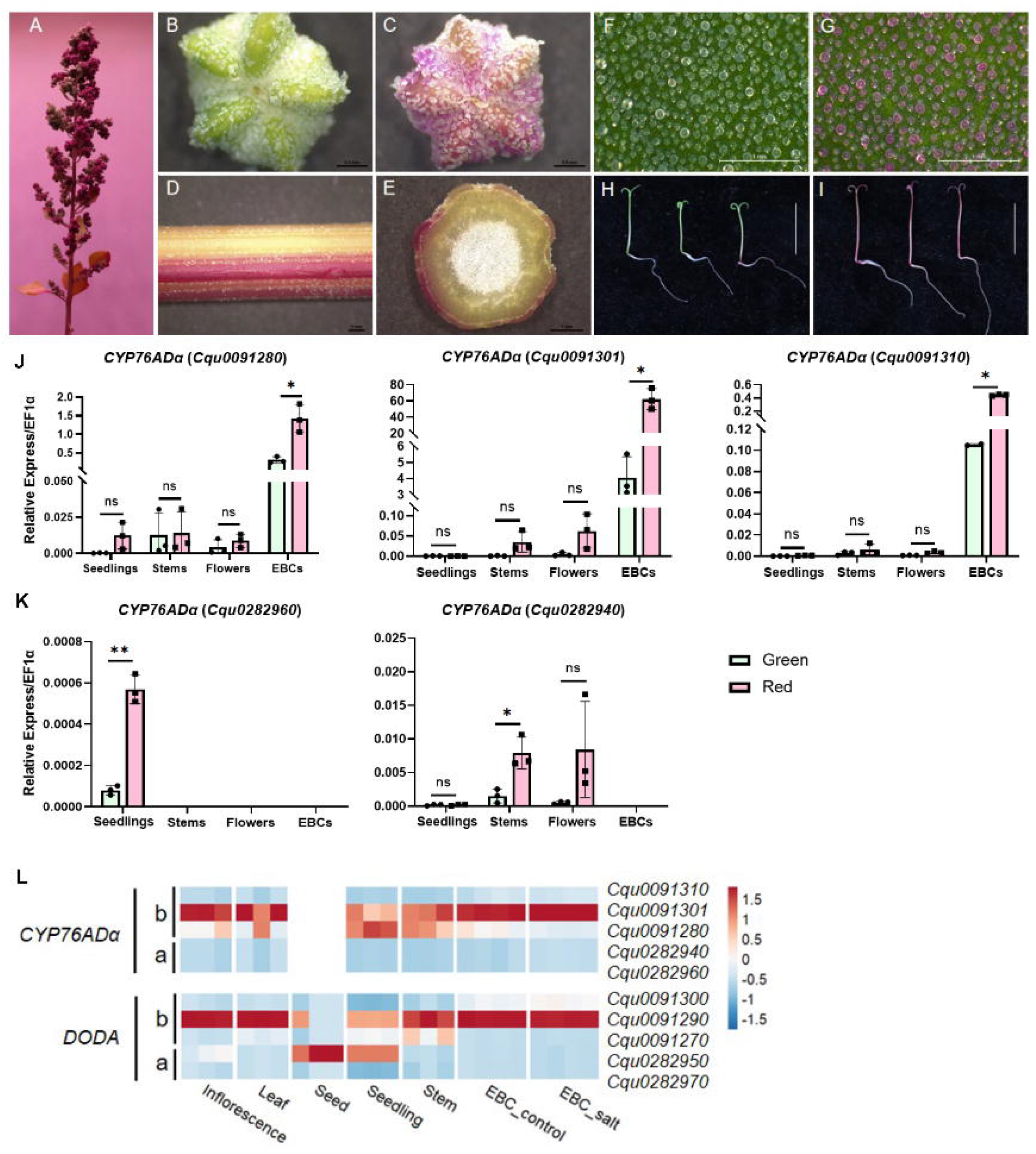

To broadly assess the contribution of the *CYP76AD*α–*DODA* gene pairs, we examined our earlier quinoa transcriptome datasets in the Real variety (Zou et al., 2017). In most tissues, except for dry seeds and seedlings, the B-subgenome genes *Cqu0091290* (DODA) and *Cqu0091301* (CYP76ADα) were the most highly expressed within their families (**Figure 4L**). Since *Cqu0091301* in the Real variety is structurally incomplete (**Figure 3A**), the primary functional contribution in these tissues (inflorescence, mature leaves, stems) is likely borne by the second-highest expressed B-subgenome pair (*Cqu0091270*-*Cqu0091280*). Thus, although the A-subgenome genes are preferentially expressed in non-EBC tissues, the B-subgenome *CYP76ADα* genes provide a consistently higher overall contribution to betalain biosynthesis across the majority of plant organs.

## Discussion

Quinoa is a globally important halophytic crop valued for its nutritional density and adaptability to harsh environmental conditions, including high salinity, drought, and low-temperature stress. Betalains, the key pigments analyzed in this study, possess significant antioxidant capacity, making them critical for mitigating stress-induced oxidative damage. Furthermore, increasing consumer demand for natural food colorants – driven by a shift away from synthetic additives – has magnified the commercial value of betalains, with red beet being the primary source (Nemzer et al., 2011; Davies, 2018). While the core enzymatic steps of the betalain biosynthetic pathway are known, the regulatory mechanisms governing this pathway are less understood compared to the extensively studied anthocyanin pathway.

In this study, we harnessed the variegated phenotype of quinoa accession P0429 to dissect the regulation of betacyanin biosynthesis. We established that the red and green mosaic pattern is fundamentally a cell-type-specific phenomenon driven by the differential accumulation of betacyanins within the epidermal bladder cells (EBCs). Specifically, spectrophotometric quantification demonstrated that red EBCs accumulated significantly higher betacyanin levels than colorless EBCs, and that the pigment difference was overwhelmingly restricted to the EBC layer, with the underlying leaf lamina contributing only marginally to the color contrast (**Figure 2**). Betalains are synthesized in the cytosol and sequestered in the vacuole (Khan, 2016); thus, these distinct accumulation patterns directly determine the resulting tissue coloration. Since EBCs represent a uniform and readily isolable cell type, the variegated leaves provided a uniquely advantageous system for transcriptome profiling, allowing us to pinpoint regulatory mechanisms at the cellular level.

Transcriptome profiling of isolated red and colorless EBCs from the same leaves yielded a highly specific set of seven differentially expressed genes (DEGs). Among them, *Cqu0091301*, encoding a CYP76ADα-type cytochrome P450, was the most highly expressed and strongly upregulated gene. The *CYP76ADα* family is known to catalyze the pivotal first committed step (hydroxylation of L-tyrosine to L-DOPA) in the betalain pathway, establishing *Cqu0091301* as the likely rate-limiting factor. Neither RNA-seq nor qPCR analyses detected significant expression differences in any other known betalain biosynthetic genes (**Figure S5**), confirming that transcriptional regulation primarily targets the initial step catalyzed by CYP76ADα. Two paralogs, *Cqu0091280* and *Cqu0091310*, also showed upregulation by qPCR, but their absolute expression levels were 1–2 orders of magnitude lower than *Cqu0091301* (**Figure 4L**), suggesting a minor functional contribution to the red phenotype.

A significant finding of this study is the discovery and validation of a structural genomic variation that restores the function of *Cqu0091301*. Reference quinoa genomes, such as Real and QQ74, contain large deletions that truncate the second exon of *Cqu0091301*, resulting in a non-functional CYP76ADα enzyme. Our resequencing of the variegated accession P0429 identified a ∼4-kb genomic insertion that corrects the deletion, yielding an intact gene capable of encoding a complete P450 domain (**Figure 3**). This demonstrates that a complete and functional *Cqu0091301* allele is a necessary prerequisite for the high-level betacyanin accumulation observed in the red EBCs of variegated varieties.

Furthermore, we found that *CYP76ADα* and *DODA* genes, which catalyze sequential steps in the pathway, are organized in adjacent pairs across the quinoa genome (**Figure 3C**). This gene clustering is frequently observed in secondary metabolic pathways (Boycheva et al., 2014) and is often associated with the coordinated regulation of enzyme production, conferring evolutionary advantages and enhanced environmental adaptability (Zhan et al., 2022). Our expression data support this concept of co-regulation: while *CYP76ADα* genes are generally expressed at an order-of-magnitude higher basal level compared to *DODAs* across various tissues, their relative expression profiles remain highly similar (**Figure 4L**), suggesting they are governed by common regulatory mechanisms.

Interestingly, we also identified a reverse-transcribed, transposon-derived gene (*Cqu0091311*), encoding a zinc-finger protein, within the intron of *Cqu0091310* in both the reference and variegation accessions. Given that *Cqu0091311* is specifically expressed and significantly upregulated in red EBCs (**Figure S1D**), it may represent a previously unknown *cis*- or *trans*-acting regulatory factor responsible for the cell-type-specific activation of the betalain pathway, a hypothesis that warrants future investigation. These observed genomic variations in the B-subgenome homologs are consistent with a recent publication suggesting that the B-subgenome is more dynamic (Jaggi et al., 2025).

We observed that the *CYP76ADα* homologs exhibit distinct and non-overlapping tissue expression preferences based on their subgenome origin. The three *CYP76ADα* copies from the B subgenome are preferentially expressed in EBCs, while the two A-subgenome copies (*Cqu0282960* and *Cqu0282940*) are silenced in EBCs but show induction in other pigmented tissues like stems, inflorescences, and seedlings (**Figure 4J, K**). This pattern clearly demonstrates functional divergence between the A and B subgenomes, where the EBCs rely mostly on B-subgenome copies for pigmentation.

The prevalence of structurally defective B-subgenome alleles, like the truncated *Cqu0091301* in reference genomes, suggests that many quinoa varieties may lack EBC pigmentation. Our overall expression analysis, however, shows that the B-subgenome homologs, especially *Cqu0091301*, provide a consistently higher total contribution to betalain biosynthesis across most mature pigmented organs (**Figure 4L**). In such varieties, betalain accumulation would primarily be mediated by the other complete *CYP76ADα* genes in non-EBC tissues. This phenomenon highlights a resilience mechanism in the allotetraploid genome, where two subgenomes have specialized, allowing different tissues to utilize different paralogs for pigmentation, thereby ensuring the maintenance of an essential stress-mitigating compound like betalain throughout the plant. These findings offer new molecular insights into the adaptive roles of subgenome divergence in allotetraploid species.

## Materials and methods

### Plant materials

The quinoa plants (accession P0429, P0556, and P0571) were grown in the growth room at the Chenshan Botanical Garden Research Center. Plants were cultivated for 40 days under a day-night cycle of 14-10 hours and 28-22ºC, with a light intensity of 300 μmol·m^-2^·s^-1^. Leaves exhibiting the variegated phenotype were sampled from quinoa plants with consistent growth. For transcriptome analyses of EBCs, each biological replicate was harvested from five individual quinoa plants.

### Extraction and measurement of betacyanins

Weigh 0.1 ∼ 0.5 g of tissue (EBCs, leaf lamina, or whole leaf) from both the red and green sectors of the variegated quinoa phenotype. Grind the samples into a fine powder in a pre-chilled mortar. Immediately transfer the pulverized sample to 7 mL of pre-cooled methanol at 4ºC. Shake vigorously and incubate for 30 minutes at 4ºC. Centrifuge the mixture at 12,000 rpm for 10 min, discard the supernatant, and retain the precipitate. Resuspend the precipitate in 3 mL of double-distilled water. After incubating for 30 min at 4ºC, centrifuge again at 12,000 rpm for 10 min and collect the aqueous supernatant. Measure the absorbance value at 538 nm using a UV spectrophotometer.

### Library preparation and sequencing

Quinoa seeds were surface sterilized and grown in ½ MS medium supplemented with 0.7% agarose in a Percival tissue culture chamber. Genomic DNA was extracted from two-week-old seedlings using DNeasy Plant Kits (Qiagen). Epidermal bladder cells were brushed off the leaf surface using cosmetic brushes directly into liquid nitrogen. EBCs were homogenized on a TissueLyser II (Qiagen), and total RNAs were extracted from the EBCs using RNeasy Plant Mini Kit (Qiagen). The genome resequencing and transcriptome libraries were prepared at the Genomics Core Facility of Shanghai Center for Plant Stress Biology, Center for Excellence in Molecular Plant Sciences, following standard protocols, and sequenced on the Illumina NovaSeq platform with paired-end 150 bp sequencing mode.

### Transcriptome analysis

The raw reads were subjected to quality control and adapter trimming using trim_galore with a quality score threshold of 20. The processed clean reads were used for *de novo* assembly using the Trinity software with default parameters (Grabherr et al., 2011). Alternatively, hisat2 (v2.1.0) was used to align the clean reads to the quinoa Cq_real_v1.5 reference genome. Picard markdupulication was then used to remove PCR duplicates. The processed alignment files were used in featureCounts (v1.6.3) to count the number of high-quality, strand-specific, and uniquely mapped reads corresponding to each gene. The resulting read counts for each annotated gene were subjected to data reproducibility testing using the edgeR package in R. The TMM algorithm was applied for between-sample normalization, and FPKM values were calculated as the expression level for each gene. The batch information was included as a variable in the linear model during the calculation of differential gene expression significance using the generalized linear model. Genes with an adjusted p-value smaller than 0.01 and over 2-fold changes after fitting to the generalized linear regression model were defined as Differentially Expressed Genes (DEGs).

### Reverse transcription and quantitative PCR

Total RNA was reverse transcribed into cDNA using the TransScript One-Step gDNA Removal and cDNA Synthesis Kit (TransGene, Beijing) for the quantification of candidate gene expression levels. The mRNA component of the extracted quinoa EBC total RNA was used as the template for reverse transcription, performed according to the manufacturer’s suggestions. Quantitative PCR was performed on a CFX96 Real-Time PCR system (Bio-Rad) using standard protocols, using primers listed in Table S1.

## Data availability

The Cq_real_v1.5 genome assembly and annotation were deposited at CoGe (http://www.genomevolution.org/) with genome ID 69570.

## Author contributions

Z.Z. and Y.W. performed experiments, analyzed data, and wrote the manuscript; X.H., T.Y., J. G., G.F., and T.Z. performed field experiments and analyzed data; H.Z. designed the project and wrote the manuscript.

## Acknowledgements

We thank all members of the Zhang Lab for helpful discussions. We thank the Core Facility for Genomics of the Shanghai Center for Plant Stress Biology (PSC) for constructing the sequencing libraries and the Core Facility for Bioinformatics of PSC for maintaining the high-performance computing (HPC) clusters.

