## Supplementary material for "Genomic Structural Variation Underlies Cell-Type-Specific Betacyanin Variegation in *Chenopodium quinoa*": Dataset S2

```
  1        10        20        30        40        50
  |        |         |         |         |         |
  ATGGATAATACAAGCCTAGCAATGATACTTGCAATTTGGTTCATTGCTTT
  TCATTTCATTAAAATATTATTTACTAGCCAAACTTCCAAACTTCTTCCTC
  CAGGCCCTAAACCACTTCCAATAATCGGCAACATTCTTGAAGTTGGTGAC
  AAACCTCACCAGTCATTTGCTAACCTCGCCAAGATTCACGGCCCTCTAAT
  ATCTCTACGTCTAGGCAGTGTCACAACTATTGTTGTATCATCAGCTGAAG
  TAGCCAAAGAAATGTTCTTAAAAAAAGACTACCCTCTTTCTAACCGTACT
  GTTCCTAATTCTGTCACTGCTGGTGACCACCACAAACTCACCATGTCGTG
  GTTGCCTGTCTCCCCAAAGTGGAGGAATTTTAGAAAGATCACAGCCGTTC
  ATTTACTTTCTCCTCAAAGACTTGATAGTTGCCAAAGCCTTAGGCATGCC
  AAGGTACAACAACTTTTTCAATATGTACAAGAATGTGCACAAAAAGGGCA
  AGCCGTTGATATTGGCAAGGCTGCATTTACTACATCCCTCAATTTGTTAT
  CAAAACTATTCTTTTCGGTCGAATTAGCCCATCATAAATCCCATACATCT
  CAACAATTCAAAGAACTTATATGGAATATTATGGAAGATATTGGCAAGCC
  TAACTATGCTGATTACTTTCCAATCTTAGGATGTGTCGATCCTTCGGGTA
  TTCGACGGCGATTAGCGTCTAGTTTTGACAAGCTAATTGCTGTTTTTCAA
  AGTATAATCACTCAAAGGCTTGGTAGTACAACAACAAAGATAAATGATGT
  GCTTGACGTTCTTCTCGACCTCTACAAACAGAAGGAGCTTAGCATGGCCG
  AGATTAACCATCTTCTAGTCGTAAGTTCATTTTTTTTCTCCATATATATA
  TATATATATATATATATATATATATATATATATATATATATATATATATA
  TATATATATATATATATTCTAATGTAATCTTATGACTTTTGACTACCACA
  ATTTTTTATAATGGATGGCTGAGATTATTTGTTTTTAGATTTATTTTTAA
  TTAGTAATTTTATTTTTAAAAAAATCATTTTTAATTATTTTATATATATG
  TTTTATTTCCTTAATTTTGAAAACATAAAAATAATTTCTGAAAGATTAAA
  ATATATATTACGTATTTAAAATGATTAAAAATTAATTTTAATCAAATTTA
  CGAATAAAAAATAAATTAAAAAATAAATAATCTCAGCCGTCCATTTTAAT
  AAAATTGTGGTAGTCTAAAGTCATAAAACTACATTGAAATCATATATATG
  TTTTCTTGTGTACGTAGTTTCGAGGAAATTGACGTAAATAGAATATCATA
  TCAGACTAGTTCTTAGATTCACGTCAAAGATTTTGACGAATTAAGCTAAT
  TACTACAATCATCAAGCATGCATCTACTAATTCTTAATATTTTTTCTTTA
  CCAATAATGAGCTTGTGCATGGCATAAAATGATTTTTTAAGAAACACAAC
  TTTACATTATAAAATTAATAACGAATTACGATTCATTGATTGGATATATA
  TATATGTGACAAATATATAAACCACATTTATTTGGAGCTTGTTAAACTCT
  GACCTGTCACATTAGATCAGAATTTAAAGTATACTTGTTACTGTTTTCAT
  CTCAAATAATTGTTTTACTATACTATGCTTTGCAACAAAAACAAGTAAAA
  TAGTATATTTGTTGATAAAAATTGTACGAGTAAGATTTTGATGTATATAA
  TGAAAAAGGCAAAATAGTATGAGAAGATAGAACGCAACCCGTGCATTCAC
  AATAATAAAACAAGAAAATTCCATTAAGAACAATTAAATGATTGGTAAGA
  ACATTATTCTATAAAATAAAGAAAGTATACATTGTTTAATAGAATAATAG
  ATACATTCTAGCTAAAAGTGGCGAACACAAAAAGAGCGATGTTTGTGATT
  ATAAGAGTGTTAGACCTCACATTTTCTACGAAAAGTGCAAGGGTGCAAAT
  GATTAGACCCCACAAATATCTACAAAGAAATTAATTAAAGAAAATTCATT
  GATTAGGTTATATTGCATTCTTGTAATAATTGATAGAATAAGTAATAAAG
  CTATTAAATTAATTATTGCCAACTTATTTCGGTTTAATTGCTTAAATTAA
  TTTAGTATAATTTATGCATGTCAGGATATATTTGATGCCGGGACTGACAC
  TACATCAAGTACTTTTGAATGGGCAATGGCAGAGTTAATTCGAAATCCTA
  AAATGATGGACAAAGCTCAAAAAGAAATTGAGCAAGTCTTGGGCAAGGAT
  AGACAAATTCAAGAATCAGACATTATTAAGTTACCTTACTTACAAGCCAT
  TATCAAAGAAACATTGCGACTACACCCACCAACTGTATTTCTCTTGCCTC
  GTAAAGCTAATTGTGATGTTGATTTATTTGGCTATGTTGTGCCAAAAGAT
  GCACAAATACTTGTTAATTTATGGGCTATCGGTAGAGATCCTCAAGCATG
  GGTGAACTCTGATGTGTTTTTACCTGAGAGGTTTTTGGGATCCGAAATTG
  ATGTAAAGGGGAGAGATTTTGGACTCTTACCTTTTGGAGCTGGAAGGAGA
  ATATGCCCAGGGATGAATTTGGCTATTAGAATGTTAACTTTGATGTTAGC
  TACGCTTCTTCAATTCTTCAATTGGAAGCTTGAAGAAGGTATGAACCCAG
  AAGATCTAGACATGGATGAAAAATTTGGAATTGCCTTACAAAAGACTAAA
  CCTCTTCAGATCATTCCGGTTCTTAGGTATTATTGATCGTTGTCAAATGT
  TTACGTATTTATATGTTTTTGTCAGTAAATTCATTCACTTTCTAATTGTT
  TAAGTTTTCTACAAATGTTCATGTTTGTTTGAATCTTCCATTCAATGCAA
  TTTTAAGGTGGTACTAGAGATCAAGTTTATGTACGTTAGCTCATATGCTA
  TTTATTGTAATCTATGTTTACATATAAATTGTTAATAAAAGTTTGTCAAT
  TGTTTGATGGATATTTTTGGTGATTTCATCAGCAGTTTTTTACTGTTCAT
  AATACATGCTATTTAGGCCAAGTCTTCATAGCAAGCTAGTGTGAAACTTC
  TATTCTAATCTCCTTGTTTTTTTAGGAAAATTTGACATTTGCTACCACCC
  AAAACGCCTCGCTTTAGATTTGCTACCATTTTAATTTTTTTTTACGTTGC
  TACCACCTAATTCAAGTTTGTTAGAAATTACTACTACTTAACGGAATCCG
  TTAGGATTTCCGTTAAGTTTATTAAAAAAAAAAGAAAAAGAAAAAAACCC
  AGCCACCCCACTTTGCCACTCTTTCCCCTCAACTGCCACCCCTCCCCCTG
  CCGGCGGCCGGCGCCGGCGACCTCACCCCCCTCCCTTTTCCCCACCCACG
  AAGGCAACCCTCCTCCCTTTTCCTCACCCATGAATCCAAATCTACCACCC
  ACGAAGCCAACCCCCACCGAGATCAGCCCCCCTCCCCGTTACCCCACCGG
  AATCGAACCCCTACCCTTCCTCCACCCAAAAACTGACACCCCTAACCCTA
  AACCACCAGCTCGAAACCGCCTAAACGGGTTTTAGGGTTTCGCTTGTGGC
  TGATTGGGGGGTGTTGCAAATTGTGGTGGCTGCAAATTGAGGAGGGTGCA
  AATTAAGGTGGTTGCGAATTGAGGAGGCTGCAAAATTGGTGGCGGAGTTG
  GTGTTGCAGGTGGGTGCGAATTGGTGGGTGCAAATTGAGGGTTTGCGAAA
  TTGGTGGCTGAGTTGGTGGTTGCGAATTGGTGGTTGATTTGGTGGAGGGA
  ACGGGGAGGGGGGGTTGAATACGGTAGGGGTTGGCTTCGGTGGGTGGGGG
  AAAGGAAGGTGGGGGTTGGCTTCGTGGTGGGGGAAAGGGAAGGGGGTTAG
  CTTCATGGGTGGGGGAAAGGGAGGGGGAGGGGTCGTCGGCGCCGACCGTA
  GTTTTGGGCGCCGACCGCTGGTAGGAGGGAGGGGGAGGGGTGGCAATTGA
  GGGGAGGGGGTGGCAGTGGGGTGGCTAGACGGTGGTACGGCCCCACTGAC
  GGAAATTGTAACGGTGTAAGGCATGGGGTAGTAATTTCTAACAAACTCTG
  TTTAGGGTGGTAACAAAGTAAAAAAAAAAAATTAAACGGTGGCAAATCTG
  AAGGTAGGTGTTTTAGGTGGTAGAAAATGTCAAAATTTCCTTTTTTTTAC
  TAGCTAGTTCAAAGGAAATTGTAAAGAATTTTAGATGGAGACTTCTATCT
  CTCGACACCACAACATGATTGTGACTTGGCTCAATGTTACTGAGAGGAAT
  TAAGCTTCAACAGAGGTCCAATACAATAAGAAAAAGCCCCATCAAGTAAT
  TTGACCTCATTTGGCATATTTTGTAGTCAATCTTAACTCAATTATTACTA
  TGTCATATTTATGATAAAATATTAGCACAACTATTAGGGTAGCTTCAACC
  TCTAGGAACTTAAGAGGCATGGCTCTAGCATAATTAAGTCCTTCCCTTAA
  TGAATAGAGTTCCGCTCTTAGAGCATTAGGAGCATTATATTTTCTAGAAT
  AGCCAAAATGCCACTCTCCTTTTGCATTACTAAAAGCACCACCTCCACCT
  CCTAATTTTGTAGAAACCCAAGCACCATCTGTGTTTAATTTCAGAAACCC
  TGGCTTTGGGGACGCCAGCTAATATCCACGCTTTTTAGAAATTTTGTAGG
  ATTCAAGGTCTGGTGTCGAAAAGACTTGAGATTATAGAGGATACATCTCG
  GGGTCATATAATTCCGTTTGGAACCATATATGATCATTGTATGTTGTCAT
  AAATTTTCGTGGTAGGCTTGATTTAATGAAATCGGTGCAAGTATTTACCA
  GTAAATTAGTCGTTTAAAGGCAAATTATATGGTTACACCCACCCAATTTG
  ACTCCCGGAGGATATTAGGGACCATGTTAAACGTTTCGGCAATTTTTATT
  TCCTGACGAATAAAAGTCAAAAATTCCAAAACTAAAGGCTAAAATACACG
  CTTTTTGACAAATTTTAGAGGTATCAAGGCCCGATGTCGAAAATCCTTGA
  AATTTTTGGGGATACATCTCGAGGTCATATAATTCCGTTTGAAACAATAT
  ATGATCATTATATATTGTCATAAATGTTCGTGGTAGGCATGATTTAATGA
  AATCGGTGCAAGTTAATTGTAAACCGGTAAATTGAACGTTTAAAGGGAAA
  TTGGATGGTTACAACCACCCAATTAGACTCCCGGAGGATATTAGGGATCA
  TGGTAAACATTTCGGCAATTTTTATTTTCGAGCAAAAAAAATGTCAAAAA
  TTTCCAAAAATGAAGACTAATATCCACGATTTTTGAGAAATTTCGAAGGT
  TTCGGGGCCCGGCGTCGAAAAGAAATGAGATTTTAGGGGATACATTGGGG
  TCATATAATGTCGTTTGAAACCATAAATGACCATTATATGTTGTCATAAA
  TGTTCGTGGTAGGCTTAGTTAATAAAGTCAGTGCAAGTTGAAAACTGGTA
  AGTTGGTCGTTTAAAGAAAAAATTAGATGGTTCAACCCTAATAATTGGAC
  TCCCGGAGGTTACTAAGGACCATTGTAAACATTTCGCCAATTTTTATTTT
  TGGGCGAAAAAAGTCAAAAAATTCCAAAATTAAAGGCTAATATCCACGTT
  TTTTAAGAAATTTTGAAGTTTTAGCGATCCGATGTCGAAAAGACTTGAGA
  TTTTAGGTAATACATTTCGGGGTCATATAACGTCATTTGGAATCATATAT
  GATAATCATTTGTTATCGTAAAAGCTCTTGGTAGGTTTGAGTTAATGAAA
  TCGGTGCAAGTTGTAAACATGTAAATTGGCCTTTTAAAGACAAATTTGAT
  GGTTCTACCCATCCAATTGGACTTCGGGAGGATATTACAGACCATGATAA
  AAATTTCGACAATTTTTGTTTCCGCACGAAAAAAAAAGTCAAAAAATTCA
  AAAATTGAACGCTAATATCCACCTTTTTGATAAATTTTGGAGCCTTTGGG
  GGGTGTCTAAAAGTCTTGAAATTCTAGGGAATACATCTTAGGGTCATATA
  ATGTCGTTTTGAAACATATGTGATCATTATTTTTGTCATAAATGTTCGTG
  GTAGGCTAGAGTTAATAAATTCGGTGCCAGTTGTCAACTTGTAAATTGGT
  CATTTAAAGGAAAATTGAATGGTTCCACCCACCCAATTGGACTCCCGGAG
  GATATTAGGGACCATGTAAAACATTTCGGCAATTTTATTTTCGGGGCCCG
  ATGTTGAAAAGAAATGAGATTTTAGGGGATACAACATCTCGGGGTCATAT
  AATTCTGTTTGAAACCATAAATGACCATTATATGTTGTCATCAATATTCG
  TGGTAGGCTTGAGTTAATGAAATCAGTGCAAGTTGTAAACCGGTAAATTG
  GTCATTTAAAGAAAAAAAATTGGATGGTTCCAACTTAGTAATTGGACTCC
  TGGAGGGTATTAAGTACCATGGTAAACATTTCACCAATTTTTATTTCTGG
  GCGAAAAAAAGTTAAAAATTTCCAATATTGAAGGCTAATATCCACGTTTT
  TTAAGAAATTTTGAAGTTTTCGGGACCTAGTGTCGAAAAGACTTGTTGTA
  GACACCCAAATGTGTCTCCCCAAAACCAAACCCAGTGATGAGTACTTTTC
  ACGTCGAAACATAATCGAATGAAGTGAAAGTGTGGAAAATCTAGTGTTGC
  GGATAATTTTAATTTCTTTTCTCTTCTCTCTGTTTTTGAAAATTTATTTT
  AAGAAAACAGCCAATCCAGTTTGGTTGGTCCATGATATGGGTCCAATATT
  AGTGTAGTTGGATGGGGCTTGCAAAGACCTTTCCAACGATATATTATAAG
  CCCAATTCCGATGAATATTGAGGAAGTTATGGCCAATTTACCAAACTGGG
  CCCGGTTTGAGCTTTATAAACGGCCCAATGATCTAGGGGAGCGCCTAATA
  TGATTTATCATTCCTAATGCAATTAGGAATGGTTTTAGTCTTCTAGATTC
```

**Dataset S2** The genomic sequence of P0429 containing the complete *Cqu0091301* gene and the ~4-kb insertion. Note: the genomic sequence is in the same direction as the open reading frame of *Cqu0091301*.

Orange-colored letters: Exon sequence of *Cqu0091301*

Grey-shaded letters: the 3,957-bp insertion sequence in P0429 and P0556
